# The Goldilocks effect of lake size on within-population diversity in stickleback

**DOI:** 10.1101/678276

**Authors:** Daniel I. Bolnick, Kimberly Ballare

## Abstract

Many generalist species consist of disparate specialized individuals, a phenomenon known as ‘individual specialization’. This within-population niche variation can stabilize population dynamics, reduce extinction risk, and alter community composition. But, we still only vaguely understand the ecological contexts that promote niche variation and its stabilizing effects. Adaptive dynamics models predict that intraspecific variation should be greater in environments with two or more equally-profitable resources, but reduced in environments dominated by one resource. Here, we confirm this prediction using a comparison of threespine stickleback in 33 lakes in on Vancouver Island, Canada. Stickleback consume a combination of benthic and limnetic invertebrates, focusing on the former in small lakes, the latter in large lakes. Intermediate-sized lakes support generalist populations, which arise via greater among-individual diet variation, not by greater individual diet breadth. These intermediate lakes exhibit correspondingly greater morphological diversity, while genomic diversity increases linearly with lake size. These results support the theoretical expectation that habitats with an intermediate ratio of resources are “just right” for promoting ecologically relevant intraspecific diversification.

## Introduction

Many animal species that appear to be ecological generalists are in fact heterogeneous assemblages of relatively specialized individuals (Bolnick *et al.* 2003; Araújo *et al.* 2011). Ecologists have therefore become increasingly interested in evaluating the community and ecosystem consequences of diet variation among co-occurring individuals (Bolnick *et al.* 2011; Des Roches *et al.* 2018). Theory and experiments have demonstrated that this within-population variation can increase population stability and reduce extinction risk (Agashe 2009), promote species co-existence (Doebeli 1997; Schreiber *et al.* 2011), change community composition (Ingram *et al.* 2011; Start & Gilbert 2017; Start 2018), and even alter ecosystem properties (Vrede *et al.* 2011). As these ecological effects depend on the magnitude of among-individual variation, we need to understand the causes of individual diet specialization, and in what settings it will be more or less pronounced. (Araújo *et al.* 2011).

The most widely accepted explanation of individual specialization invokes frequency-dependent selection arising from resource competition (e.g., Levene 1953; Wilson & Turelli 1986; Doebeli 1996b). Consider a consumer population inhabiting an environment with two functionally distinct resources. The consumer might specialize entirely on whichever resource is most profitable (taking into account nutritional value and abundance, Stephens & Krebs 1986). But, as this preferred resource becomes scarcer due to exploitation, the newly more-abundant under-used resource becomes relatively profitable (Bolnick 2001). Consequently, the consumer should evolve an intermediate phenotype that uses both resources (even if modest trade-offs penalize generalists). In population genetic models of a diploid organism, this leads to a balanced polymorphism dominated by heterozygotes that may or may not be especially well suited to either resource (Levene 1964; Hedrick 1986; Wilson & Turelli 1986). In quantitative genetic models, the population trait mean converges towards an intermediate generalist (Doebeli 1996a; Schreiber *et al.* 2011). If trade-offs limit efficient use of both resources, that generalist may experience persistent disruptive selection that increases trait variation and individual specialization (Nuismer *et al.* 2005). Thus, the equilibrium trait mean and variance should both depend on the relative availability of alternative resources.

This diversification process is thought to be especially relevant when populations invade new environments. In such settings, the invading population can be released from the constraints of interspecific competition (Costa *et al.* 2008; Bolnick *et al.* 2010), permitting niche expansion. Meanwhile, intraspecific competition can be strong, which acts to favor individuals who adopt new kinds of resources and thereby mitigate competition with their own species. The ‘niche variation hypothesis’ (NVH) posits that this niche expansion arises not via greater individual niche breadt, but via increased among-individual variation (Van Valen 1965; Bolnick *et al.* 2007). Experiments have confirmed this: increased intraspecific resource competition drives disruptive selection (Swanson *et al.* 2003; Bolnick 2004; Svanbäck *et al.* 2008) that promotes increased diet variation among individuals (Svanbäck & Bolnick 2005, 2007a). Some other comparative and experimental studies have challenged this NVH model (e.g., Parent *et al.* 2014; Jones & Post 2016).

So far, most studies seeking to explain variation in individual specialization have focused on the effects of competition (reviewed in Araújo *et al.* 2011). But, a growing number of comparative studies have found that resource diversity (often termed ‘ecological opportunity’) leads to greater population niche breadth via increased individual specialization (Parent & Crespi 2009; Martin & Pfennig 2010; Araújo & Costa-Pereira 2013; Evangelista *et al.* 2014; Cloyed & Eason 2016; Yurkowski *et al.* 2016; Costa-Pereira *et al.* 2017), consistent with the NVH (Van Valen 1965). Most tests of the NVH have focused solely on either patterns of diet variation (Bolnick *et al.* 2007), or morphological variation as a proxy for diet variance (Rothstein 1973; Patterson 1983; Meiri *et al.* 2005). To date, no studies have simultaneously evaluated the effects of resource diversity on among-individual diet variation, morphological variance, and genetic variation.

This study tests the hypothesis that resource diversity promotes individual specialization and greater trait diversity in a single consumer species. The simplistic example described above, with a balance between two resources, makes a very specific prediction. When the ratio of two resources changes along an environmental gradient (or differs among habitat patches), there will be an intermediate point along the gradient where the resources are equally profitable (considering abundance and nutritional value), maximizing ecological opportunity. We therefore expect a consumer population’s niche breadth, and individual specialization, to be greatest at these intermediate points along the gradient; a “Goldilocks effect” (referencing the children’s story “Goldilocks and the Three Bears”, in which a child enters the home of a family of bears and searches for porridge that is “just right”, neither too hot nor too cold).

To test for a Goldilocks effect promoting intraspecific ecological diversity, we use lake fish that balance benthic and limnetic resources. Around the world, lake fish species have evolved to either specialize on large benthic prey on the lake substrate or on limnetic mid-water zooplankton, or use a mixture of both (Moodie & Reimchen 1976; Lavin & McPhail 1985; Robinson & Wilson 1994; Kusche *et al.* 2014). Benthic prey tend to be relatively more abundant in small lakes dominated by shallow littoral habitat. Limnetic prey dominate in large lakes, where volumes become large relative to the shallow perimeters that support benthic prey. Therefore we expect that resource diversity (the evenness of benthic and limnetic prey) is low in small and large lakes, but maximized in intermediate sized lakes (Fig. 1). Consistent with this expectation, disruptive selection on trophic morphology is strongest in threespine stickleback populations inhabiting intermediate-sized lakes (Bolnick & Lau 2008). As a corollary, we expect individual specialization to be greatest in those intermediate-sized lakes as well. Because individual specialization reflects underlying diversity in morphology (Snowberg *et al.* 2015), one might expect morphological trait variance to also be highest in these intermediate lakes (although Nosil and Reimchen (2005) argued for a positive linear trend with lake size. Using a comparative study of lake populations of threespine stickleback (*Gasterosteus aculeatus*), this paper confirms the “Goldilocks effect”: individual specialization is greatest in intermediate-sized lakes.

**Figure 1.**
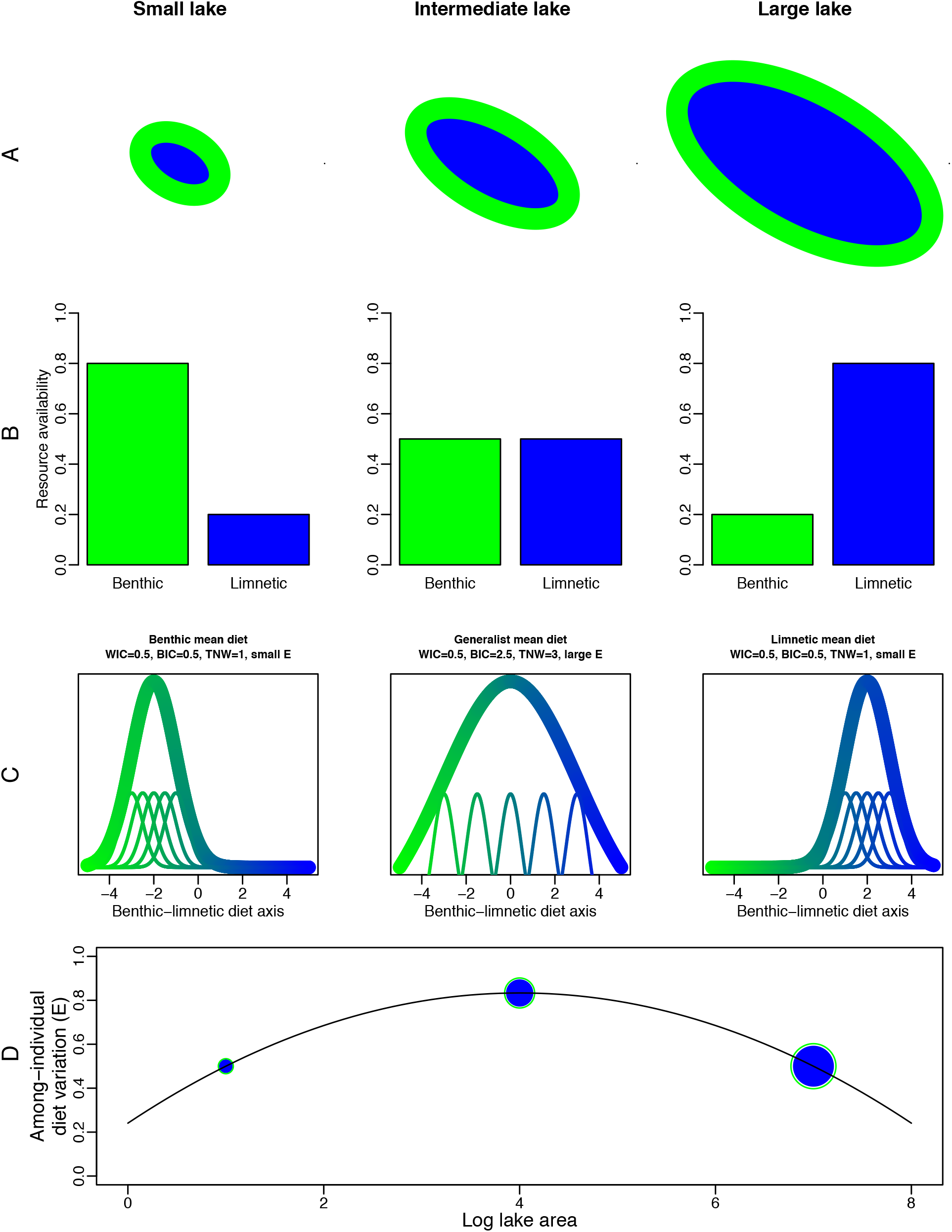
A schematic diagram of the hypothesis tested in this study. Row A: three lakes of increasing size, with shoreline benthic habitat (green) and mid-water limnetic habitat (blue). Row B: the corresponding resource availabilities of the three lakes, with decreasing relative abundance of benthic prey in larger lakes. Row C: Because of the resource availability shifts in Row B, the populations’ diets shift from benthic (in a small lake) to limnetic (in the large lake). The diet distribution of the population as a whole is represented by the thick taller line, and is made of up diets of individuals (smaller lines, color coded to represent relative use of benthic and limnetic prey). Row D: As a result of the shifting diet distributions in Row C, among-individual diet variation is expected to be greatest in the intermediate-sized lake.

## Methods

In June 2009 we collected between 60 and 100 threespine stickleback from 33 lakes on Vancouver Island, British Columbia, spanning a range of lake sizes. Fish were captured in unbaited minnow traps set for less than three hours, and immediately euthanized and preserved in formalin. Collection and animal handling were approved by the University of Texas IACUC (Protocol # 07-032201), and a Scientific Fish Collection Permit from the Ministry of the Environment of British Columbia (NA07-32612).

All fish were weighed, measured for standard length, and sex determined via dissection. A random subset of 30 fish per lake were also measured for gape width, gill raker number, and gill raker length. Gape width and gill raker length were size-adjusted by calculating residuals of log transformed values regressed on log length. We calculated trait means and variances for each lake. These data also form the basis of two other studies focused on parasite metacommunity structure, and methods are explained in greater detail in Bolnick et al. (BioRxiv, a,b).

For a random subset of 23 lakes we also enumerated stomach contents on the same subset of 30 fish. We recorded presence or absence of prey taxa in each fish’s stomach. Stomach contents are an admittedly coarse and cross-sectional sample of individuals’ diet, in stickleback reflecting approximately the previous 6 hours of foraging (Svanbäck & Bolnick 2007). However, previous work has demonstrated that stomach content variation among individuals is a robust measure of diet variation in the population, being correlated with individual morphology, and with variance in stable isotope signatures that reflect long-term diet over months (Matthews *et al.* 2010; Snowberg *et al.* 2015). Direct observation of foraging individuals also confirms there is variation in foraging microhabitat (benthos versus limnetic zone), which is correlated with those individuals’ stomach contents and stable isotopes (Snowberg *et al.* 2015).

We categorized prey as benthic or limnetic, and, per fish, calculated the proportion of present prey taxa that were benthic. Non-metric multidimensional scaling analysis yielded a first major axis that was tightly correlated with the proportion benthic prey, so we used the latter, more intuitive, metric. The total number of prey taxa observed per fish provides a metric of diet richness, with the recognition that this is a brief cross-sectional sample (for discussion of such caveats, see Bolnick *et al.* 2002; Araújo *et al.* 2007; Araújo *et al.* 2011). We calculated individual specialization using the metric *E*, which measures among-individual diet disparity. *E* ranges from 0 when there is complete diet overlap between individuals, to 1.0 when every individual uses unique resources with no overlap with other individuals (Araújo *et al.* 2008). This is simply 1.0 minus the mean pairwise diet overlap (*IS;* Bolnick et al. 2002). We calculated *E* using RInSp (Zaccarelli *et al.* 2013).

We sampled fin clips from each fish before preservation in formalin, and extracted DNA from a random subsample of 12 fish per population. We genotyped 175,350 single nucleotide polymorphisms (SNPs) from 336 fish (107,698 SNPs scored per fish on average), using ddRADseq (Peterson *et al.* 2012). Lab and bioinformatic protocols are detailed in (Stuart *et al.* 2017). We calculated genome-wide heterozygosity for each fish and then averaged these to obtain the average heterozygosity for each lake.

### Statistical Analyses

The focal hypothesis of this study is that individual specialization (measured by the metric *E*) will be maximized in intermediate-sized lakes, mid-way along the benthic-limnetic diet continuum. We first tested the assumption that lake size is associated with population mean diet, by linear regression of the mean proportion benthic prey as a function of log lake area. Having confirmed this linear trend, we next ran a quadratic regression of diet variation, *E*, as a function of log lake area, anticipating a negative quadratic gradient. To confirm this, we also used quadratic regression relating *E* to mean proportion benthic prey, and a larger model with linear and quadratic effects of both lake area and mean proportion benthic prey.

Increased population niche breadth could instead arise via increased individual niche breadth. To test this possibility, we used quadratic regression to test the relationship between individual diet breadth (prey richness) and either log lake area, or mean proportion benthic prey. The NVH predicts this relationship to be flat.

To test the role of morphological variation in diet diversification, we calculated the standard deviation of each morphological trait (standard length, gill raker number, size-adjusted gill raker length, and size-adjusted gill raker number). We used multiple regression to test whether *E* increases with these traits’ standard deviations. We then used quadratic regression to test whether each trait standard deviation is highest in intermediate-sized lakes.

Most genetic variation is expected to be approximately neutral, and so genomic diversity (mean heterozygosity) should be associated with population size rather than ecological or morphological variation. We therefore used linear regression to test for a positive relationship between mean heterozygosity and log lake area, whereas we expected no relationship between mean heterozygosity and *E*, or trait standard deviations. In contrast, loci involved in adaptation to benthic or limnetic environments should show allele frequency correlations with lake size. If these allele frequencies span from near 0 to near 1 across the range of lake sizes, then polymorphism should be greatest in intermediate sized lakes. To test this prediction, we iterated through SNPs, focusing on loci genotyped in at least 50 individuals and with minor allele frequencies exceeding 0.1 in the entire dataset (to ensure reasonable power and minimize multiple test corrections). For each SNP we used a binomial general linear model to regress allele frequency (out of the number of genotyped individuals in each population) as a function of log lake area. We also tested for correlations between SNP allele frequency variance, and diet variation (*E*).

## Results

As commonly assumed, stickleback in larger lakes tended to consume relatively more limnetic than benthic prey (Fig. 2A; linear regression log lake effect P < 0.0001; all regression results summarized in Table 1). This trend confirms past studies (Lavin and McPhail, 1986). Because the populations range from 10% to 90% benthic prey, intermediate populations are indeed ecological generalists that use roughly equal mixtures of benthic and limnetic resources.

**Figure 2.**
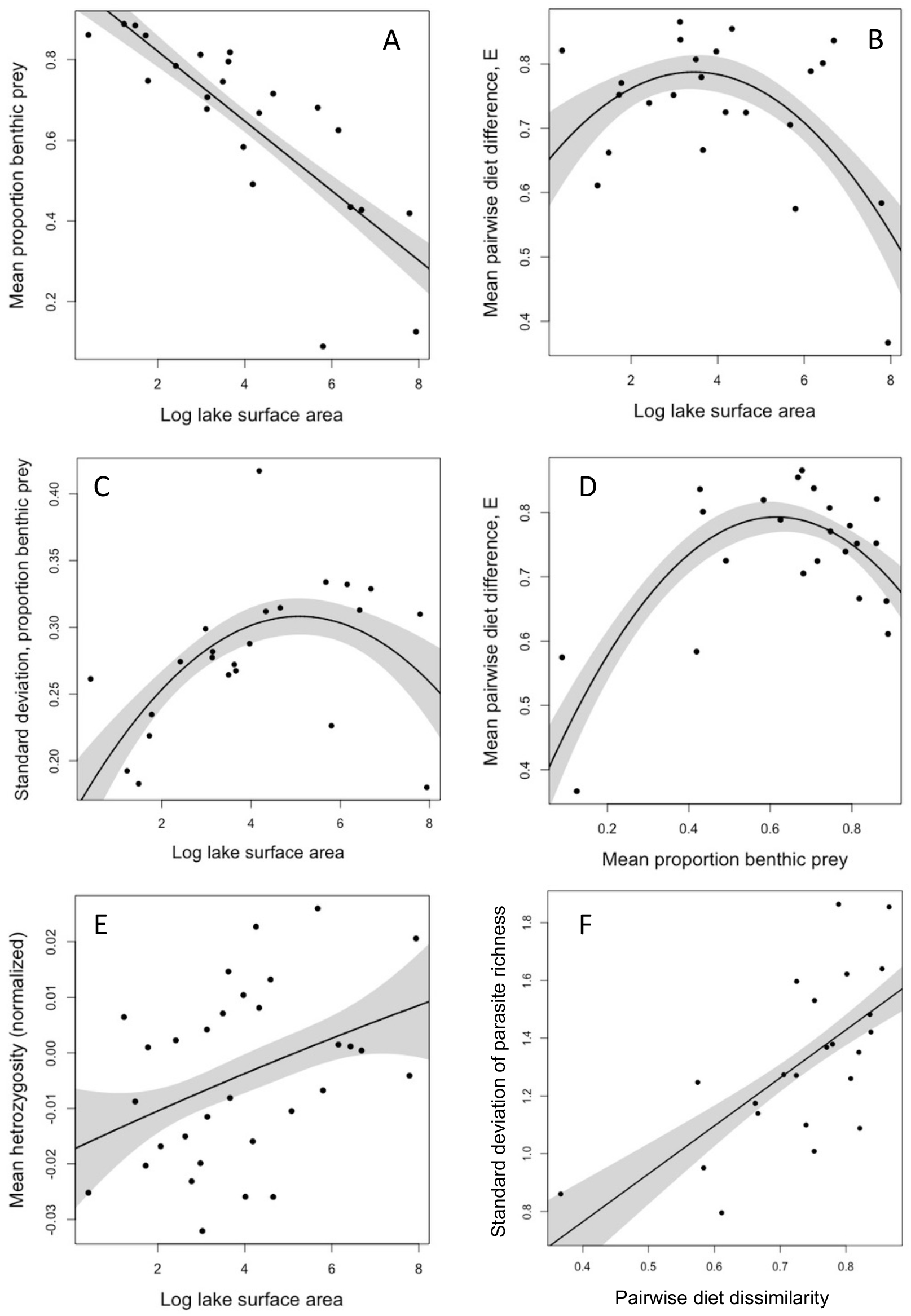
Linear and quadratic regressions examining predicted relationships between population mean diet, diet variation, genetic diversity, parasite diversity, and lake area (log hectares). Points represent lakes as the level of replication. Lines are linear or quadratic regression estimates, shaded regions are one standard error confidence intervals. Statistical support for the trends reported here are provided in Table 1.

**Table 1.** Results of linear and quadratic regressions between focal variables. Correlation tests (where the direction of causation is ambiguous) are reported only in the main text. Some models test *a priori* linear predictions, so quadratic effects are omitted. For each model, the table lists the figure where the relationship is ploted, the linear (and where relevant quadratic) slope estimate, its standard error, and a P-value testing the null hypothesis of zero slope, and for each model there is an r^2^ value provided as well. NA denotes relationships not plotted as a figure.

| Figure | Dependent | Independent | Linear effect | Linear se | Linear P | Quadratic effect | Quadratic se | Quadratic P | $r^2$ |
| --- | --- | --- | --- | --- | --- | --- | --- | --- | --- |
| 1A | Mean proportion benthic prey | Log lake surface area | **-0.086** | **0.013** | **<0.0001** | - | - | - | 0.66 |
| 1B | E | Log lake surface area | **0.084** | **0.040** | **0.0481** | **-0.012** | **0.004** | **0.0140** | 0.36 |
| 1C | SD proportion benthic prey | Log lake surface area | **0.058** | **0.019** | **0.0066** | **-0.006** | **0.002** | **0.0159** | 0.35 |
| 1D | E | Mean proportion benthic prey | **1.540** | **0.325** | **0.0001** | **-1.25** | **0.31** | **0.0005** | 0.58 |
| 1E | Heterozygosity | Log lake surface area | **0.0003** | **0.001** | **0.0297** | $4.9 \times 10^{-5}$ | $6.5 \times 10^{-4}$ | 0.939 | 0.15 |
| 1F | SD per fish parasite richness | E | **1.662** | **0.406** | **0.0005** | - | - | - | 0.44 |
| S1 | Individual diet breadth | Log lake surface area | 0.056 | 0.132 | 0.677 | -0.006 | 0.015 | 0.651 | 0.01 |
| S2 | Individual diet breadth | E | 0.221 | 0.581 | 0.707 | - | - | - | 0.01 |
| NA | Individual diet breadth | Mean proportion benthic prey | -0.245 | 1.336 | 0.857 | 0.184 | 1.258 | 0.885 | 0.00 |
| S3 | E | heterozygosity | -1.456 | 1.79 | 0.425 | - | - | - | 0.03 |
| S5 | SD standard length | Log lake surface area | 1.42 | 0.716 | 0.0567 | **-0.197** | **0.081** | **0.0217** | 0.21 |
| S6 | SD size-adjusted gape width | Log lake surface area | 0.015 | 0.008 | 0.0675 | **-0.002** | **0.001** | **0.0495** | 0.13 |
| NA | SD size-adjusted gill raker length | Log lake surface area | 0.051 | 0.067 | 0.451 | -0.009 | 0.007 | 0.241 | 0.11 |
| NA | SD gill raker number | Log lake surface area | 0.005 | 0.100 | 0.963 | 0.003 | 0.011 | 0.777 | 0.05 |

Individual specialization was most pronounced (highest *E* values) in intermediate-sized lakes, as predicted (Fig. 2B. quadratic effect P = 0.014, Table 1). As a follow-up confirmation, we calculated the among-individual standard deviation in their proportion benthic prey for each lake, which is also quadratically related to lake area (P = 0.0159, Table 1, Fig. 2C). This trend exists because mid-sized lakes contained stickleback with an intermediate diet (Fig. 2A), and individual specialization (*E*) is strongest in populations with an intermediate diet (Fig. 2D). When we shifted to a multiple regression to simultaneously consider how individual specialization (*E*) depends on quadratic effects of lake size and benthic diet, we found statistical support only for the latter effect (diet P = 0.0002, diet^2^ P = 0.0001, area P = 0.945, area^2^ P = 0.188, model r^2^ = 0.727). Thus, intermediate-sized lakes have intermediate-diet stickleback, which promotes greater individual specialization.

In contrast, individual diet breadth does not contribute to the trends described above. There is no detectable correlation (linear or quadratic) between individual niche breadth and log lake size, or with *E* (both P >0.5; Table 1, Figs. S1 & S2). This result thus corroborates the central tenet of the Niche Variation Hypothesis, that generalist populations arise via increased among-individual variation, while individual niche breadth remains unchanged.

There was no linear correlation between individual specialization (*E*) and genome-wide heterozygosity (r = −0.174, P = 0.425; Fig. S3). However, heterozygosity was greater in larger lakes, as would be expected with greater effective population sizes (Fig. 2E; P = 0.030, Table 1). For 2149 of the 41,284 SNPs examined, allele frequency was correlated with log lake size, including 5 loci that survived Bonferroni correction (Fig. S4). None of these alleles exhibited significant watershed covariate effects. Some of the loci exhibited a wide range of allele frequencies, ranging from 0 (fixed with the reference genome nucleotide) to 1.0 (fixed for the derived allele) across the range of lake sizes. Such SNPs were most polymorphic in intermediate-sized lakes. The strongest association was found in two neighboring SNPs on linkage group 4 (bases 19204072 and 19204307, P = 0.0000096 and 0.000068, respectively), which are within 100 kb of four genes, *foxp2*, *gpr85*, *tmem168a* and *ifrd1*. A second site, on linkage group 20 (site 15807116, P = 0.000074) lies between the genes *bola1* and *nr2f5*, and within 100 kb of *sv2a*. A site on linkage group 1 (site 7503783, P = 0.000092) sits between *cluha* and *tlcd2*.

There was no linear correlation between individual specialization and the standard deviation of any single phenotypic trait (standard length, r = 0.178, P = 0.414; size-adjusted gape width r = 0.327, P = 0.128; size-adjusted gill raker length r = 0.137, P = 0.532; gill raker number r = −0.004, P = 0.985). However, some traits were more variable in intermediate-sized lakes: both standard deviation body length (Fig. S5) and size-adjusted gape width (Fig. S6) were greater in mid-sized lakes. However, the standard deviations of gill raker number and size-adjusted gill raker length were unrelated to lake area (Table 1). Some of this trait diversity was, surprisingly, negatively correlated with mean heterozygosity (gill raker length r = −0.402, P = 0.0278), while other traits were marginally correlated with heterozygosity (standard length r = −0.335, P = 0.066) or uncorrelated (gape width r = 0.106, P = 0.576; gill raker number r = −0.082, P = 0.666). The most noteworthy correlation between individual specialization and population phenotypes actually involved parasitism. As described elsewhere (Bolnick et al, BioRxiv a,b), we also enumerated parasite infection loads for all sampled fish. Populations with greater individual specialization exhibited greater among-individual variation in per-fish parasite richness (Fig. 2F, r= 0.666, P = 0.0005).

## Discussion

Many theoretical models suggest that increasing resource diversity can lead to the evolution of a polymorphic generalist consumer (Levene 1953; Wilson & Turelli 1986; Rueffler *et al.* 2006). The ecological generalization may arise via greater among-individual variation rather than greater individual niche width (Van Valen 1965). Our results confirm this expectation, providing clear observational evidence that resource diversity (‘ecological opportunity’) promotes within-population variation in diet and morphology but not neutral genomic variation.

Like many lake fish, stickleback consum both benthic and limnetic resources (Lavin & McPhail 1985; Lavin & McPhail 1986). The ratio of these resources is dictated by the ratio of the lake perimeter to open water, which increases with lake size. We confirmed Lavin and McPhail’s (1985) finding that stickleback diet changes linearly with log lake area: small lakes contain predominantly benthic-feeding stickleback, and large lakes contain mainly limnetic-feeding stickleback.

At some intermediate lake area, the ratio of benthic to limnetic prey must be roughly balanced. In the absence of trade-offs, a consumer might evolve a generalist strategy in which all individuals use both prey. But, we do not see a correspondingly higher individual diet breadth in the intermediate-sized lakes where stickleback have a generalist diet. This observation is consistent with previous experiments which found that individual niche breadth was relatively insensitive to inter- and intraspecific competition (Svanbäck & Bolnick 2007b; Araújo *et al.* 2008; Bolnick *et al.* 2010). Instead, stickleback in intermediate-sized lakes are more likely to experience disruptive selection on trophic morphology (Bolnick and Lau 2008), which should promote diet diversity (Svanback and Bolnick 2007). Accordingly, the present results demonstrate that the generalist stickleback in these lakes achieve their broader ecological niche via greater among-individual diet variation and morphological variation. Variation in body size and gape width is correspondingly higher in intermediate-sized lakes as well, consistent with a previous study reporting that morphological and dietary variance within populations are positively correlated (Snowberg *et al.* 2015). However, this result contrasts with Nosil and Reimchen (2005), who found a positive relationship between trait variance and lake size in other stickleback populations from smaller islands in coastal British Columbia. They used lake volume rather than area, so we cannot directly compare our results, but it appears that their survey sampled just the smaller half of the quadratic trend we examine here; very large lakes are not found on the islands they surveyed. Their positive trend may thus be reconciled with the results here, whose slope is positive in the smaller half of the lakes studied.

An alternative explanation is that diet variation is effectively neutral, arising simply from neutral genetic diversity. This hypothesis is not supported, as genome-wide genetic diversity increases linearly with lake size. This positive trend is expected as larger lakes should contain larger effective population sizes, and matches previous results from microsatellites (Caldera & Bolnick 2008). Consistent with the mostly neutral behavior of most genomic markers, genome-wide diversity is unrelated to phenotypic or diet variation.

Although genome-wide heterozygosity should be roughly neutral, there were some apparently non-neutral loci whose allele frequency was strongly correlated with lake area. SNPs whose frequencies spanned from 0 to 1.0 across the range of lake sizes were most polymorphic in intermediate-sized lakes. These loci therefore exhibit positive correlations between their genetic diversity and the degree of individual specialization. Whether these loci have direct phenotypic effects on diet, or confer adaptations to other aspects of lake size, is unclear. It is noteworthy that the genomic locus most strongly linked to lake size (on LG4) contains multiple genes potentially involved in learning and behavior. Of these, *foxp2* is best known for its role in language and brain development (Enard *et al.* 2002). *gpr85* is also associated with brain size (Matsumoto *et al.* 2008), and *tmem168a* is a newly discovered gene possibly linked with behavior (Fu *et al.* 2017). This locus is also close to *ifrd1* (interferon-related developmental regulator 1) which regulates both neutrophil effector function in immune response, and skeletal muscle differentiation and regeneration (Gu *et al.* 2009). The locus on LG20 is close to genes involved in protection against oxidative stress (*bola1*, (Qin *et al.* 2015)) and development of the vertebrate jaw (*nr2f5* (Barske *et al.* 2018)). Any of these genes would require extensive follow-up study with experimental genetics to evaluate their potential phenotypic effects and adaptive value. For now, we make two observations. First, genome-wide heterozygosity is unrelated to individual specialization. Secondly, there are putatively non-neutral loci whose functions are plausibly associated with habitat adaptation, and whose polymorphism is greatest in the most diet-variable populations. This raises the enticing, previously-unreported possibility of eventually finding genetic variants associated with increases in among-individual diet variation in natural populations.

Individual-level diet variation within stickleback populations can have appreciable community-wide effects. Co-occurring individuals with different diets will experience different levels of intraspecific competition (Bolnick 2004). Individuals also have different overlap with other species of fish such as trout and sculpin, common intraguild predators on stickleback (Bolnick *et al.* 2010). Here, we found that populations with greater individual specialization also exhibited greater among-individual variation in parasite richness. Many stickleback parasites are trophically transmitted, so such a connection between diet and infection disparity is to be expected. This trend (plotted in Fig. 2F) illustrates a broader point that the degree of diet variation among individual stickleback can expand to affect the entire community in which they are embedded. Ingram et al (Ingram *et al.* 2011) manipulated stickleback body size variance while keeping mean size constant, in cages in a natural lake. The degree of diet variation differed among cages, and was correlated with shifts in benthic and limnetic invertebrate abundance and community structure, indicating that diet variation within one consumer species has community-wide effects. Other studies in stickleback generated even greater trait variation by mixing together divergent populations (lake and stream, or benthic and limnetic species and their hybrids), and also found dramatic shifts in prey community structure and ecosystem properties (Harmon *et al.* 2009; Matthews *et al.* 2016).

Population genetics (Levene 1953; Wilson & Turelli 1986), adaptive dynamics (Doebeli 1996b; Ackermann & Doebeli 2004), and quantitative genetic eco-evolutionary models (Schreiber *et al.* 2011) all suggest that resource diversity can promote within-population variation, in the form of polymorphism, adaptive branching, and disruptive selection. A simple corollary is that when there exists a gradient in the ratio of two resources (e.g., benthic:limnetic availability), individual specialization should be greatest in the middle of the gradient, where the resources are most evenly balanced. The results presented here represent the first test of this theory, confirming that individual specialization (and some facets of morphological variation) are greatest in intermediate-sized lakes where stickleback populations are generalists using both benthic and limnetic prey. The implication of this finding is that certain geographic settings are more favorable to resource polymorphism and perhaps even adaptive speciation. This fits into a broader emerging literature supporting the notion that ecological opportunity promotes variability within populations (Parent & Crespi 2009; Martin & Pfennig 2010; Araújo & Costa-Pereira 2013; Evangelista *et al.* 2014; Cloyed & Eason 2016; Yurkowski *et al.* 2016; Costa-Pereira *et al.* 2017). These shifts in individual specialization should have cascading effects on prey, competitor, and parasite community structure (Des Roches *et al.* 2018), most pronounced (in this instance) in intermediate-sized lakes. However, we do not know yet whether these community effects then reciprocate by changing patterns of disruptive selection, which would represent an eco-evolutionary feedback loop mediated by changes in variance, rather than changes in trait means. Such variance-mediated eco-evolutionary dynamics are not as well understood as their mean-mediated counterparts (Hendry 2017).

## Acknowledgements

Data collection was assisted by Chris Harrison, Todasporn Rodbumrung, Travis Ingram, and Julie Day. This research was supported by a Howard Hughes Medical Institute Early Career Scientist fellowship to DIB, NSF grant DEB-1144773, and NIH 1R01AI123659-01A1.

---

**Supplemental Figure S1.**
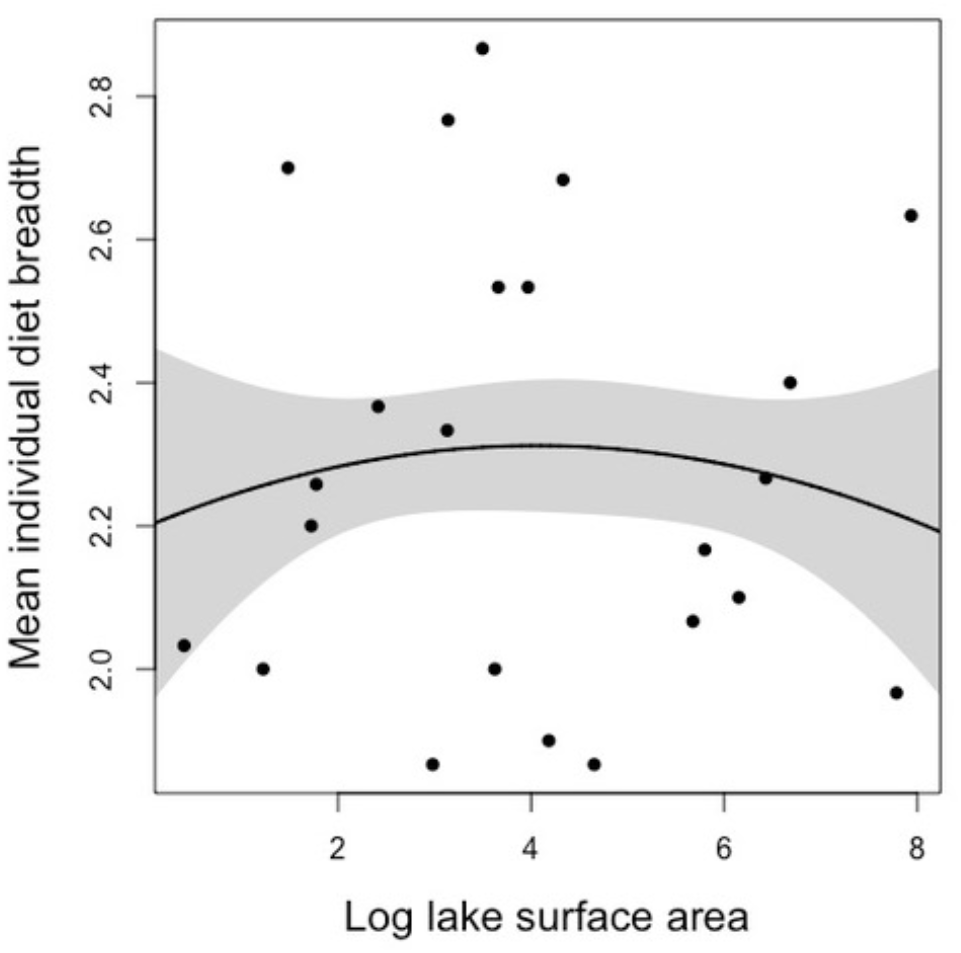
There is no significant linear or quadratic relationship between individual diet breadth and lake size. A quadratic relationship is expected if population niche expansion in intermediate-sized lakes is achieved by increased individual niche breadth (e.g., in intermediate lakes all individuals are generalists that use both limnetic and benthic prey). Statistical results in Table 1. The line is a quadratic regression estimate, the shaded region represents a one standard error confidence interval.

**Supplemental Figure S2.**
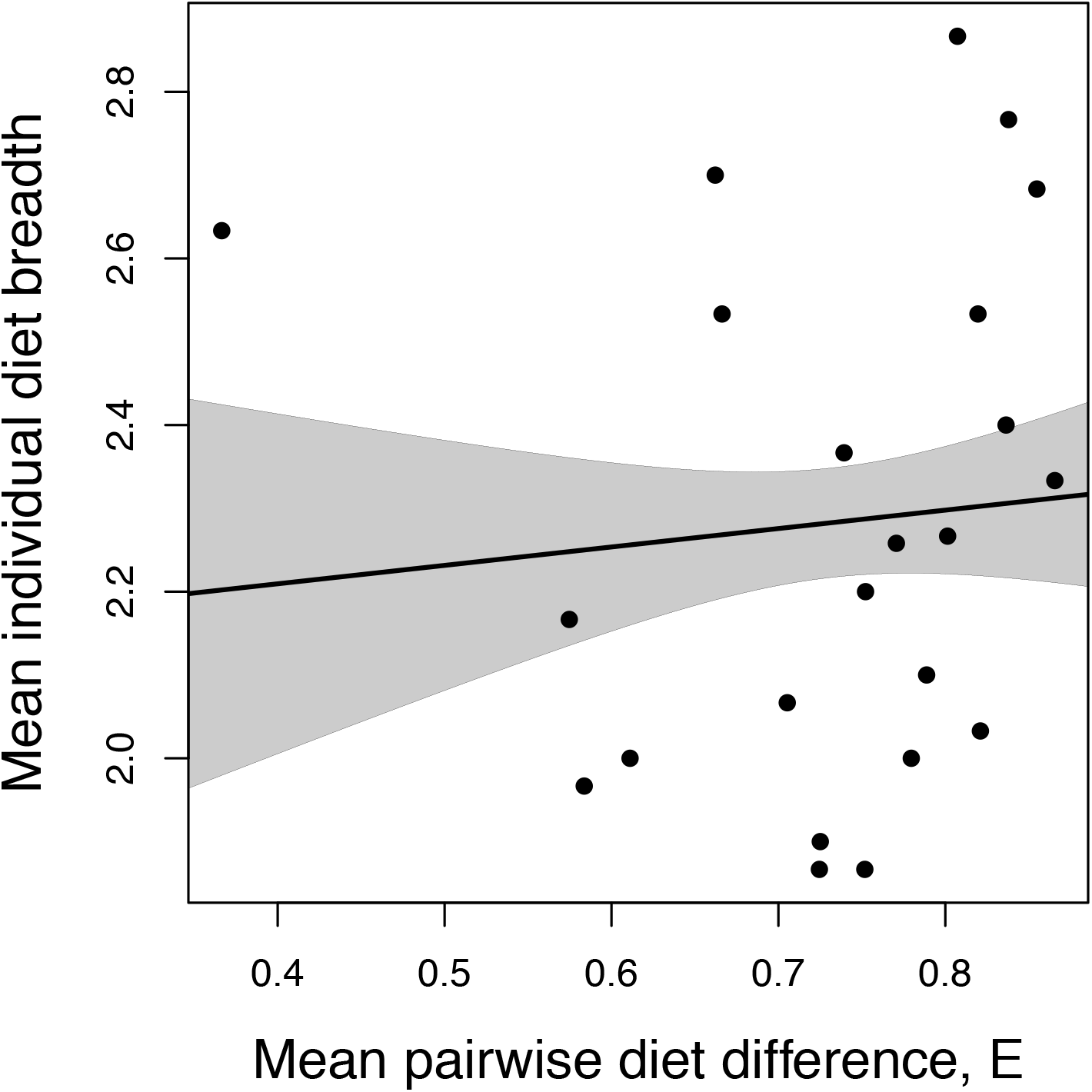
There is no significant linear relationship between individual diet breadth and the degree of individual specialization (*E*). A negative linear relationship is expected if among-individual diet variation occurs via a decrease in individual niche breadth, while between-individual differences remain constant. The lack of negative trend suggests that individual specialization arises by divergence among individuals, rather than a narrowing of individual niches. Statistical results in Table 1. The line is a linear regression estimate, the shaded region represents a one standard error confidence interval.

**Supplemental Figure S3.**
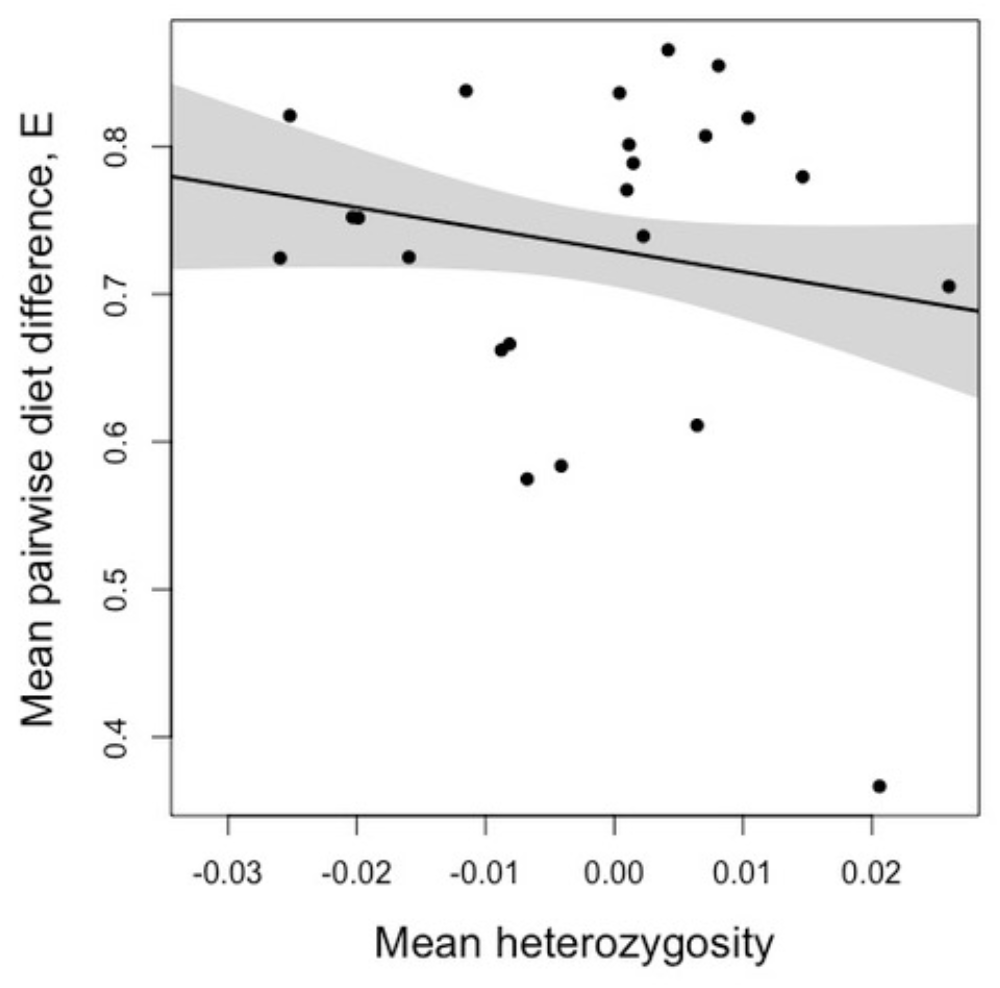
There is no significant linear or quadratic relationship between diet variation and genome-wide genetic diversity. A positive relationship might be expected if niche variation promoted genetic diversity even at neutral loci (for instance by permitting persistently higher population density), but in general we expect no effect on neutral genetic variation. Statistical results in Table 1. The line is a linear regression estimate, the shaded region represents a one standard error confidence interval.

**Supplemental Figure S4.**
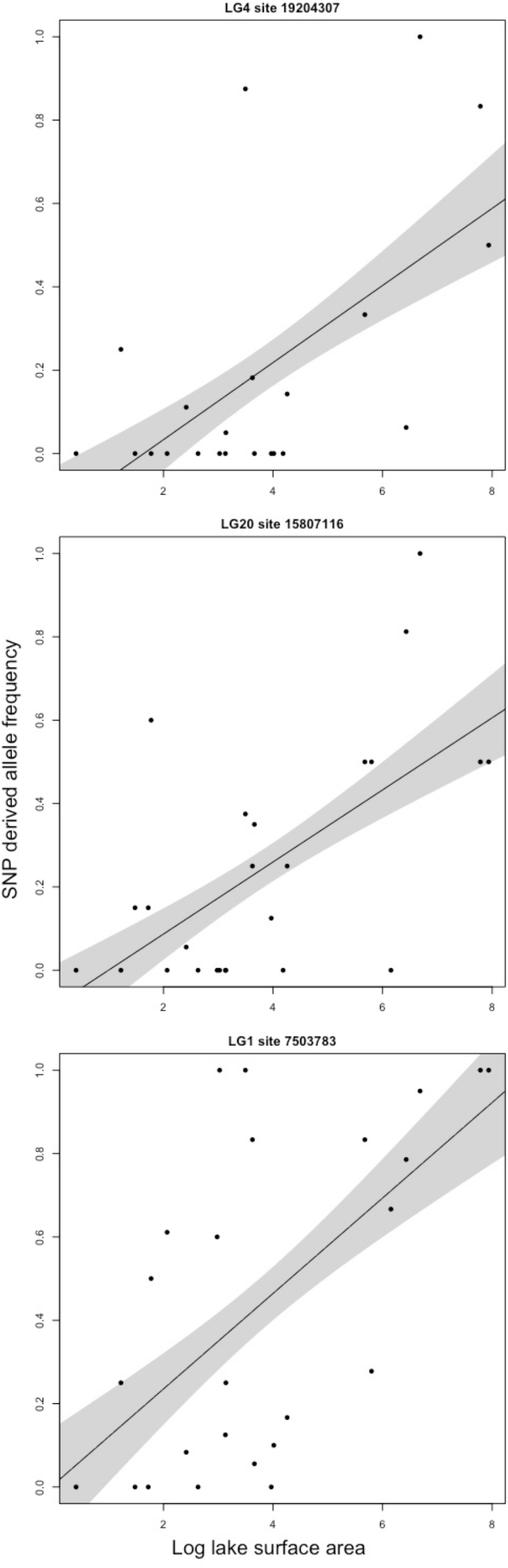
The three strongest associations between derived allele frequency (relative to reference genome) and log lake area. Points represent estimated allele frequency, the trendline and shaded region represent a binomial general linear model estimate with 95% confidence interval. The focal SNP linkage group and position is listed above each figure panel. Statistical results are provided in Supplemental Table S1.

**Supplemental Figure S5.**
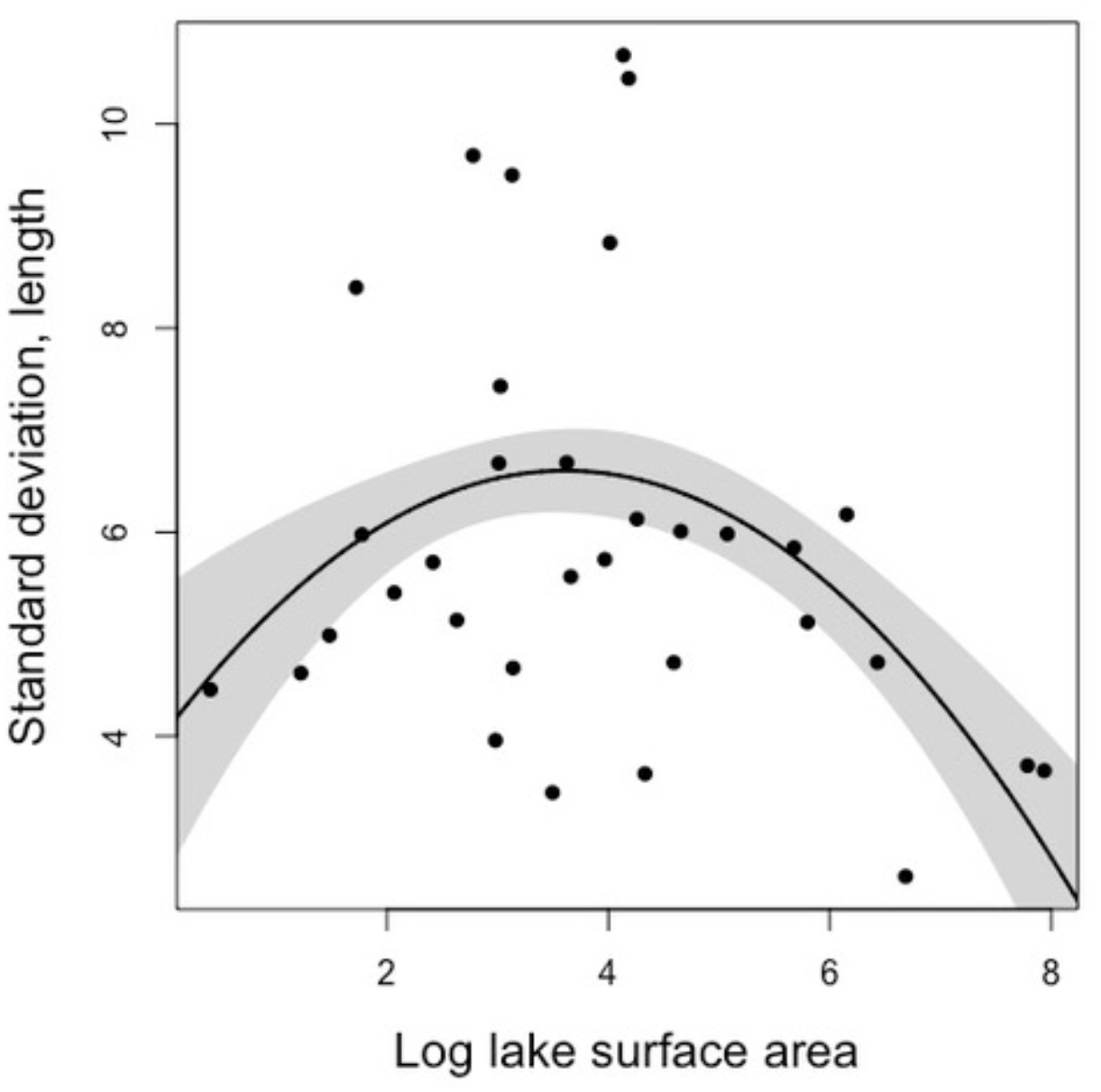
Intermediate-sized lakes support stickleback populations with greater size variation. Statistical results in Table 1. The line is a quadratic regression estimate, the shaded region represents a one standard error confidence interval.

**Supplemental Figure S6.**
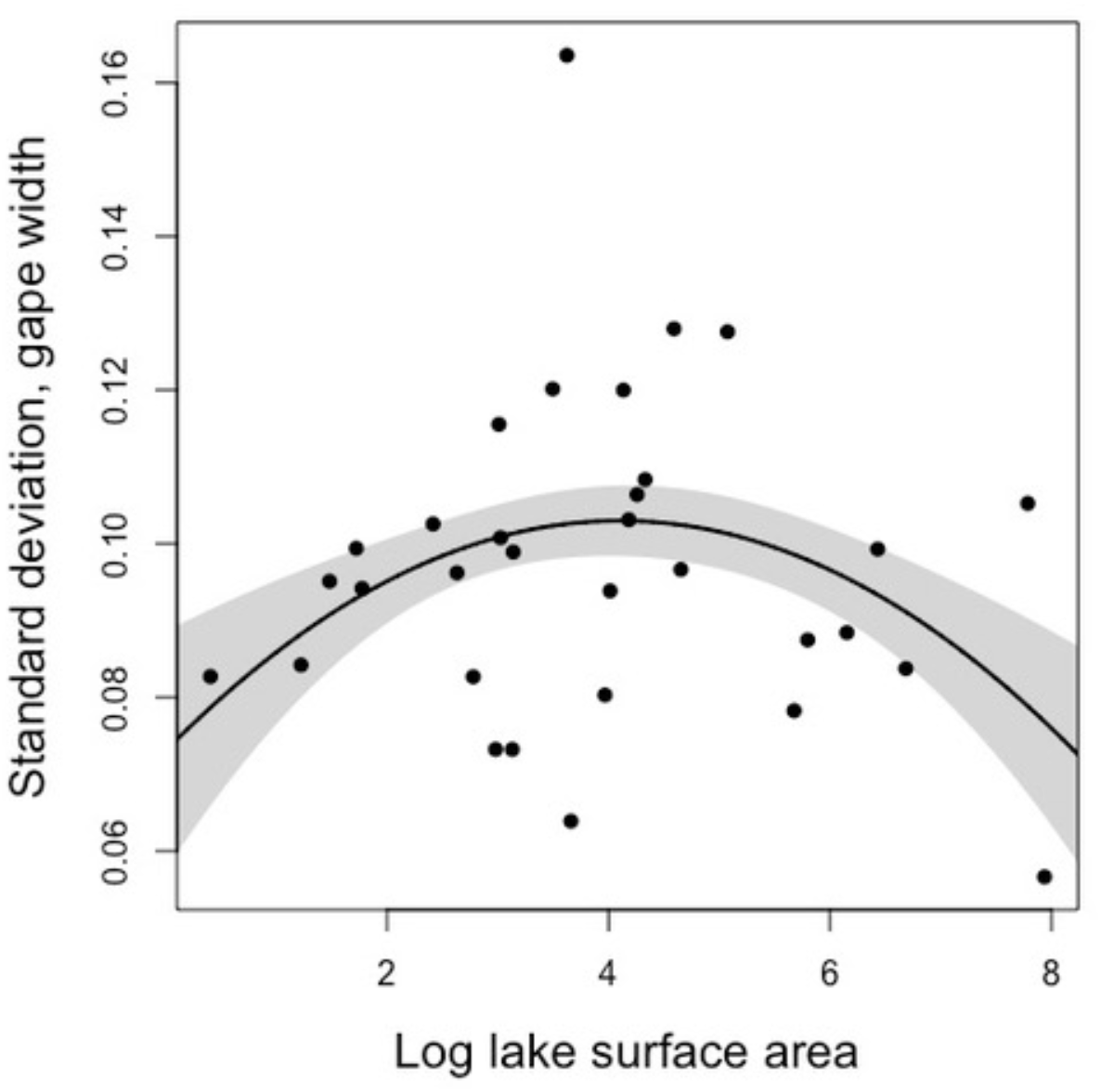
Intermediate-sized lakes support stickleback populations with greater variation in size-adjusted gape width. Statistical results in Table 1. The line is a quadratic regression estimate, the shaded region represents a one standard error confidence interval.

